# HMGB1 B-box domain self-complexes promote protein-polyelectrolyte interactions

**DOI:** 10.1101/2025.03.28.645909

**Authors:** Marten Kagelmacher, Marina Pigaleva, Ricardo Zarate, Leïla Bechtella, Kevin Pagel, Beate Koksch, Jens Dernedde, Andreas Hermann, Thomas Risse

## Abstract

HMGB1, a nuclear DNA-binding protein, can be secreted by activated immune cells or passively released from damaged cells. In such cases, HMGB1 functions as an alarmin that activates the immune system. Excessive inflammation may lead to pathogenesis, whereas this response can be dampened by polyanion binding, which impedes further receptor recognition. Moreover, HMGB1 is known to form liquid droplets in the cellular environment—a phase separation directly linked to its proper function. While the A-Box domain is believed to be primarily responsible for heparin binding due to its conserved binding site, the association and phase separation behavior of HMGB1 may be mediated by the B-box domain, owing to its extended hydrophobic regions. In this study, we first demonstrated that the B-box protein forms 30-nm large self-associates while maintaining its structure. Next, using molecularly sensitive EPR spectroscopy, we showed that the presence of these protein associations significantly enhances interactions with heparin. Notably, the local conformational changes induced by heparin are similar in both individual protein chains and their self-associated forms. To explain this effect, AlphaFold modeling was employed, revealing that the formation of protein multimers induces charge redistribution, resulting in an extended positively charged region that enhances electrostatic attraction to negatively charged polyanions, such as heparin.

## Introduction

The high mobility group box 1 (HMGB1) is a non-histone chromosome binding protein present in the nucleus of healthy cells which consists of three domains referred to as A-box, B-box, and C-terminal tail, respectively (see Scheme 1). In the nucleus HMGB1 is associated with DNA and is involved in multiple processes, like regulation of transcription, DNA repair or telomere homeostasis. Furthermore, HMGB1 can act as damage-associated molecular pattern (DAMP), if secreted from cells due to a trauma, stress, infection or cell death [^1^]. Apart from interaction with different immune receptors, it was found that its interaction with polyanions such heparin sulfate - a linear highly negatively charged polysaccharide, consisting of repeating 1-4-linked pyranosyluronic acid and glucosamine disaccharide units with an average of 2.6 sulfate groups per unit [^2^] can play an important role for the immunological response, too. To this end it has been reported that heparin sulfate can help treating sepsis as it inhibits heparinase activity and HMGB1 mediated LPS cellular uptake [^3^]. Recently, an octadecasaccharide of heparin sulfate turned out to mitigate inflammatory damage in sepsis by targeting several mediators including HMGB1 [^4^]. Therefore, it is not surprising that HMGB1 which can be considered prototypic DAMP gained interest as a therapeutic target [^5^].

As the function of HMGB1 in the biological context is associated with the binding to negatively charged polyanions, the A- and B-box domain of HMGB1, which are rich in positively charged amino acids, are considered crucial for these functions. In contrast to that, the unstructured acidic C-terminal tail (amino acid 185-215), which contains a stretch of glutamic and aspartic acids, is believed to modulate the electrostatic interaction with the polyanions. While both A- and B-box are positively charged and both are thus expected to exhibit attractive interactions with polyanions, it is reported in literature that the A-Box domain of HMGB1 is responsible for the interactions with heparin via conserved heparin-binding site (aa. 6-12), whereas the B-box plays a major role in the processes related to proinflammatory cytokine production [^6^]. It should be noted that despite such reports detailed biophysical investigations concerning these aspects are rather scarce.

The formation of functional protein multimers is a well-established phenomenon in all forms of living matter [^7^]. It is found for interfacial processes such as receptor recognition as well as in solution. A prominent example for the latter are proteins known to self-assemble abruptly to form stable aggregates (e.g., amyloid fibril formation or prion aggregation) [^8^]. Another form which has gained considerable interest in recent years is self-assemble into dynamic liquid droplets as there are evidence that such phase separations are associated with functional properties of the involved species [^9^]. In this respect, Mensah et al. [^10^] recently demonstrated that HMGB1 tends to form droplets in crowded cell environments which the authors concluded to be crucial for the protein proper functioning of the protein in DNA transcription. The latter conclusion was based on the finding that severe genetic diseases could be correlated to frameshift mutations in HMGB1 which result in a loss of charged residues on the C-terminal tail. Interestingly, these mutations disrupt the ability of the proteins to undergo phase separation providing experimental evidence for the abovementioned correlation and indicate that the formation of such condensates can be correlated with a proper function of the protein. In this work, we aim to explore the impact of such association processes on the functional properties on a molecular level. To this end we use the B-box domain of HMGB1, which possesses some hydrophobic patches along the chain and thus tend to form aggregates, to investigate differences in its interaction with heparin being the polyelectrolyte known to play an important role in the immune response. In particular, we will compare the non-associated, monomeric form of the protein with larger associated protein complexes in which the molecules exhibit very similar structural properties with respect to their ability of polyelectrolyte binding.

**Scheme 1.**
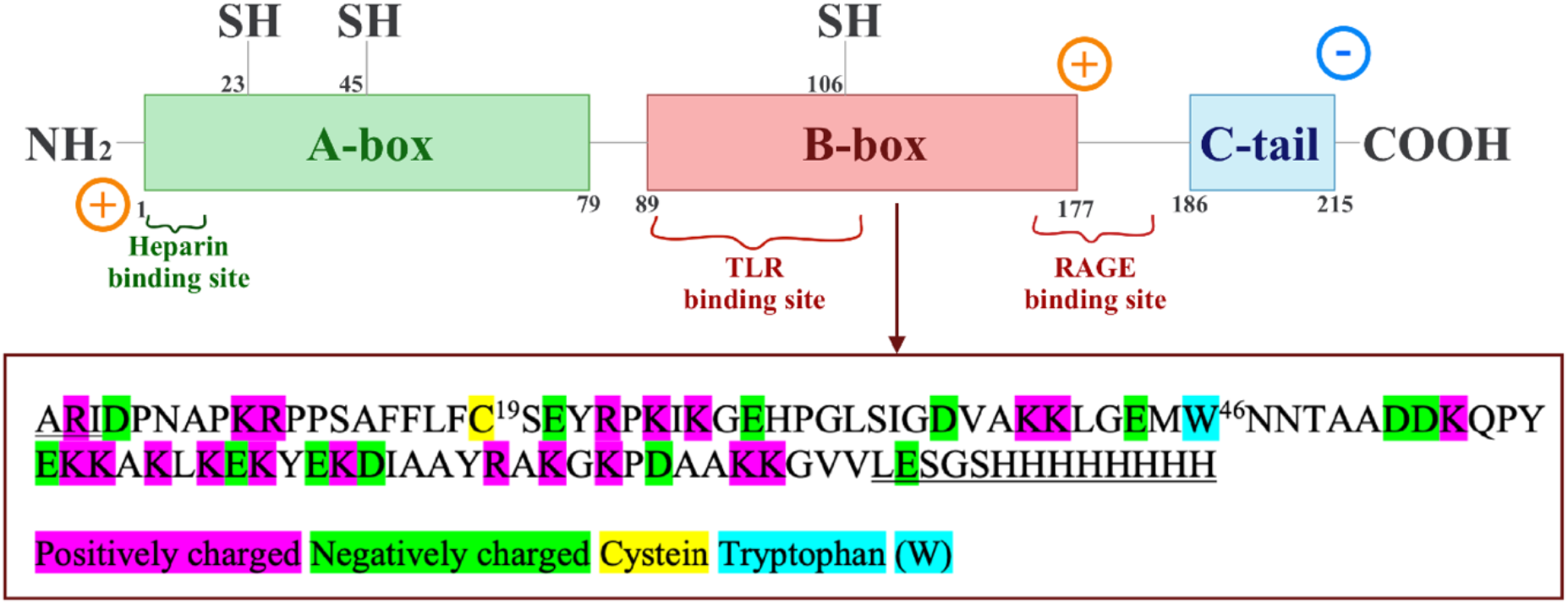
Schematic representation of the HMGB1 protein and the amino acid sequence of the B-box domain. Amino acids alanine-arginine-isoleucine- (ARI) at the N-terminus and a C-terminal 8 x Histidine (His)-tag are artificial amino acids, introduced in order to obtain a stable expression and enable purification.

## Experimental

### Cloning of the B-box domain into a modified pet22b(+) vector

The standard pet22b (+) vector (Merck, Germany) was modified by including a Serine-Glycine-Serine (SGS) and two additional histidine upstream of the His_6_-Tag resulting in a more flexible and affine 8 x His-Tag at the C-terminus. By using the following primer pair: 5’-AGCGGCAGCCATCATCACCACCACCACCACCAC-3’ and 5’-CTCGAGTGCGGCCGCAAGCTTGTCGAC-3’ in a back-to-back primer strategy the DNA was inserted. For the PCR reaction the Q5 High Fidelity 2x Master Mix from New England Biolabs (NEB, USA) was applied according the to the manufacturer’s instructions^11^. PCR Thermocycle settings were: initial denaturation 98°C for 30 s, 35 cycles with 98°C denaturation for 10 s, annealing at 50°C for 30 s, extension at 72°C for 30 s and final extension at 72°C for 120 s using the PCR 2720 Thermo Cycler (Applied Biosystems, USA). PCR products were analyzed on a 1% agarose gel and respective bands at ∼ 5.5 kb were cut out and purified with a Zymoclean Gel DNA Recovery Kit (Zymo Research, Germany). Next the purified PCR product was treated with the KLD (Kinase, Ligase *DpnI*) mix (New England Biolabs (NEB), USA) as described in the manufacturer protocol. 5 µl of the reaction mixture were used for transformation of *E. coli* DH5α cells (NEB, USA). Sequence integrity of the obtained clones were confirmed by sequencing at Eurofins Genomics, Germany. The cDNA of HMGB1 obtained from Origin, USA (pCMV6 HMGB1, NM_002129) was used as a template for cloning of the B-box domain. The following primer were used for the PCR amplification: 5’-CTTTAAGAAGGAGATATA<u>CAT</u>*<u>ATG</u>GCGCGTATT*GATCCCAATGCACCCAAGAGG-3’ and 5’-GTGATGATGGCTGCCGCT<u>CTCGAG</u>GACAACTCCCTTTTTTGCTGC-3’. The resulting PCR product was analyzed on a 1.5 % agarose gel and purified as described above. The cloning was done with the exonuclease and ligation-independent cloning (ELIC) method. [1]. In brief, the modified pet22b(+) vector was digested with the enzymes *Nde*I*/Xho*I (Thermo Fisher Scientific, USA) to create homologues ends. After gel extraction and DNA cleanup a 1:3 molar ratio (vector:insert) was used for ligation at room temperature for 15 minutes and finally transformed into *E. coli*. Insert integrity was confirmed by sequencing and restriction analysis. For better expression of the recombinant B-box domain, the DNA coding sequence of the amino acid triplet alanine-arginine-isoleucine (ARI) was inserted as part of the forward primer at the 3’ end after the methionine at the protein N-terminus.

### B-box HMGB1 domain expression and purification

The *E. coli* strain BL21(DE3) (Sigma Aldrich (USA) was used to express the protein according to a procedure described in detail by Bianchi *et al*. [^12^]. In brief, a freshly transformed colony was used to inoculate 15 ml of Luria broth (LB) medium supplemented with 100 µg/ml ampicillin (amp, Carl Roth, Germany) and 0.4 % glucose (Carl Roth, Germany). After incubation of the overnight bacteria culture at 37°C at 150 rpm, the suspension was diluted in 150 ml LB medium, 100 µg/ml amp to an initial OD600 of 0.1. After the OD600 has reached 0.7, the protein expression was induced by the addition of 150 µl of an 0.5 M isopropyl β-D-thiogalactoside IPTG, Genaxxon bioscience, Germany) stock solution, to obtain a final concentration of 0.5 mM IPTG. After 100 min at 37°C the bacteria culture was harvested by centrifugation at 8,200 x g for 30 min at 4°C.

The bacteria pellet was resuspended in 7.5 ml of lysis buffer (50 mM Tris-HCl, pH 8.0, 100 mM NaCl, 2.5 mM MgCl_2_, 10 mM imidazole), supplemented with 50 U/ml Benzonase nuclease (Merck, Germany), one spatula lysozyme (Roche Diagnostics, Switzerland) and 1 x Complete EDTA-free protease inhibitor cocktail (Sigma-Aldrich, USA). After 30 min incubation at 37°C, the bacteria suspension was lysed by ultrasonication (Sonifier 250, Branson, USA), 15% output, 30 s pulse, 1 min resting time for 15 min and subsequently centrifuged (30 min, 20,000 x g, 4 °C). After centrifugation, the supernatant was loaded onto 2 ml (equals 1 column volume (CV)) of HisPur Ni-NTA agarose (Thermo Fisher Scientific, USA) equilibrated beforehand with 10 CV of lysis buffer. After subsequent washing with 10 column volumes (CV) 50 mM Tris-HCl pH 8.0, 1.5 M NaCL, 20 mM imidazole and 10 CV 50 mM Tris-HCl pH 8.0, 50 mM NaCL, 20 mM imidazole, bound proteins were eluted with 5 CV 50 mM Tris-HCl pH 8.0, 50 mM NaCl, 500 mM imidazole.

The Ni-NTA eluted fraction was then applied to 1 ml (equals 1 CV) of a heparin sepharose 6 Fast-Flow column (GE Healthcare, USA) equilibrated with 10 CV 50 mM Tris-HCl pH 8.0, 50 mM NaCl, 500 mM imidazole. The column was washed with 10 CV of 50 mM Tris-HCl pH 8.0, 100 mM NaCl and elution of bound proteins was achieved by applying a high buffer containing a high salt concentration (2 M NaCl, 50 mM Tris-HCl pH 8.0). The eluate was further dialyzed against 50 mM Tris-HCl pH 8.0, 150 mM NaCl via repeated centrifugation in Amicon filter units (cutoff 3 kDa) (Merck, Germany). Protein concentration was finally measured at A_280_ with a NanoDrop One (Thermo Fisher Scientific, USA) using the following settings: molecular weight: 11460.03 Da and extinction coefficient: 11,460 M^-1^cm^-1^ [^13^]. The protein could be achieved in high yields of 3.15 mg/L bacterial culture and high purity > 95% as seen by SDS-PAGE.

### Sodium dodecyl sulfate-polyacrylamide gel electrophoresis (SDS-PAGE)

For protein visualization, a 15% SDS-PAGE was prepared, as described by Laemmli *et*.*al*. and Groth *et al*. [^14, 15^]. Protein samples were prepared in a 4 x Laemmli buffer with 5% β-mercaptoethanol heated to 95 °C for 5 min before loading. The gels were run for 20 min at 15 mA and 45 min at 20 mA. Afterwards the gels were stained with 40% ethanol, 10% acetic acid, and 0.2% (v/v) Coomassie G250 for 15 min. For destaining the gels were immersed into a mixture containing 20% methanol and 10% acetic acid for 15 min. For conservation, the gels were transferred to a cellophane foil from Carl-Roth, Germany and dried with a gel dryer from Bio-Rad, USA. The dried gels were scanned using a photocopier.

### Western blot

Purified recombinant B-box was further analyzed by Western blot. In brief, 50 ng of protein was separated on a 15% SDS-PAGE gel and transferred onto an Amersham Hybon-ECL nitrocellulose membrane (GE Healthcare, USA). The transfer was done in a Mini-Trans Blot Cell (Bio-Rad, USA) in a Tris/Glycine buffer (25 mM Tris-HCl, pH 8.5, 190 mM glycine) at 250 mA for 60 min, at 4°C. The membrane was then blocked with 2% BSA in TBS-T buffer (50 mM Tris-HCl pH 7.5, 150 mM NaCl, 0.1 % Tween 20) for 1 h at room temperature and shaking. 1 µg/ml of polyclonal rabbit primary antibody (detecting aa. 150-215, Abcam, United Kingdom) was then incubated overnight at 4°C with shaking. After 3 times washing for 5 min with TBS-T buffer, a 1:2000 dilution of a secondary antibody (polyclonal goat anti-rabbit labeled with horseradish peroxidase (HRP) antibody, Agilent Dako, USA) was added to the membrane and incubated for 1 h at room temperature. After subsequent washing, the dried membrane was incubated for 2 min at room temperature with an enhanced chemiluminescence solution (90 mM Tris-HCl pH 8.6, 1.26 mM luminol, 0.6 mM hydroxycoumarinacid, and 3% H_2_O_2_). Pictures were obtained with an Amersham hyper film ECL high-performance chemiluminescence film from GE Healthcare, USA using an OPTImax type TR developer (MS Laborgeräte, Germany).

### Gel Electrophoretic Mobility shift assay

To further demonstrate B-box binding to heparin in solution, fluorescein isothiocyanate (FITC) labeled heparin (Nanocs, USA) was incubated with the B-box protein and analyzed on a 12% native PA-gel. The native PA-gel was prepared as described above, except the addition of SDS and β-mercaptoethanol and without incubation at 95°C. 10 µg of both heparin-FITC and B-box protein were incubated for 15 min at room temperature in 15 µl of 50 mM Tris-HCl pH 8.0, 150 mM NaCl. 5 µl of 4 x native sample buffer was added to the mixture to result in a 1 x concentrated solution. The samples were then loaded onto the native PA-gel and run for 20 min 15 mA, followed by 40 min 20 mA at 4°C. The gels were kept in the dark during the whole procedure. Images were taken with the Versa Doc Imaging System 400 MP (Bio-Rad, USA) using an exposure time of 53 s (fluorescence: excitation 501 nm, emission 523 nm).

### Protein Filtration

To separate the single B-box domain proteins from their multimeric assemblies in the solution, 450µl of a 147µM solution of the expressed and purified B-box in EPR buffer (25 mM Tris-HCl, pH 6.9, 150 mM NaCl), was filtered through an Amicon filter with 30kDa cutoff membrane. The filters were washed with Milli Q Water, at 4°C, 14000 rpm for 10 minutes prior use in order to clean the membranes from possible cellulose residues. After that, the protein solution was placed inside the filter and spun down at 4°C and 14,000 rpm for 5 minutes, resulting in two separate solutions: one on top of the filter, where 24 µL of solution containing molecular moieties larger than 30 kDa remains, and one at the bottom, where the flow-through from the 30 kDa membrane is collected. The concentrations of the initial solution, the bottom and the top solutions from the filter were obtained using a NanoDrop spectrophotometer (Thermo Scientific, Germany) by measuring the UV-Vis intensity at a wavelength of 280 nm.

### Dynamic light scattering (DLS)

To measure the hydrodynamic radius of the initial and filtered B-box protein, DLS in backscattering mode was used on the Prometheus Panta spectrophotometer (NT.48 Series, NanoTemper, Germany). For each measurement 3 capillaries (Grade High Sensitivity 200 counts, NanoTemper, Germany) were filled with 10 µl of B-box protein solution (30 uM concentration for the initial solution and 16 uM for the filtered solution in EPR Buffer). The data was obtained at 25°C, with a DLS laser power setting of 100%. For each capillary 10 runs were performed and averaged.

### Fluorescence spectroscopy

Fluorescence spectra were taken at 25 °C after adding 100µl of a 5µM solution of the B-box protein in EPR Buffer into a Hellma micro cuvette (105.250-QS). The spectra were taken in a FL 6500 spectrofluorometer (PerkinElmer, US) using a fixed excitation wavelength of 280 nm (absorption of Trp). The emission spectrum was recorded from 300 to 400 nm, using slits of 5 nm in both the excitation and emission paths. The spectra were taken at a scan speed of 240 nm/min and a photomultiplier voltage of 500 V. The spectra shown were corrected for contributions from the buffer (see SI, Figure S1) by subtracting the corresponding spectrum.

### CD spectroscopy

200 µL of the protein solution with a concentration of at least 10 µM in EPR buffer was placed in a 1 mm light path quartz flat cuvette and measured in J-810 Spectropolarimeter (Jasco, Germany) or in DSM 20 circular dichroism spectrometer (Olis, USA). Three spectra with 301 data points each were averaged. The spectra were measured in the 190–250 nm range at a scan speed of 100 nm per minute and normalized to account for concentration and the number of residues, thereby ensuring consistency with literature data, according to equation:

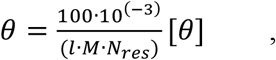

with *θ* being the molar ellipticity, *l* the light pathway length, *M* is the concentration in mol/l and N_res_ is the number of residues, [*θ*] – measured ellipticity.

### Ion mobility-mass spectrometry analysis

For the sample preparation spin labeled and unlabeled B-box HMGB1 (not filtered) were diluted to 2 µM in 100 mM ammonium acetate. Ion mobility-mass spectrometry experiments were performed on a timsTOF Pro spectrometer (Bruker, Bremen, Germany), equipped with an in-house 3D-printed offline nano-ESI source, whose design has been published [^16^]. For each measurement, 5 μL of the prepared sample were introduced in a Pt/Pd-coated glass capillary emitter prepared in-house and ionized in positive ion mode nano-ESI. The instrument parameters were set as follows: capillary voltage 1.5 kV, end plate offset -0.5 kV, dry source temperature 30°C, D1 = -20 V, D2 = -120 V, D3 = 250 V, D4 = 100 V, D5 = 0 V, and D6 = 100 V. The mass range was set at *m/z* 500-3000. The trapped IMS separation was performed in nitrogen (N_2_). Ions reversed mobility 1/*K*_*0*_ was scanned between 0.8 V·s/cm2 and 1.8 V·s/cm^2^. The accumulation time and the ramp time were set to 1000 ms and 1000 ms, respectively. The transfer time was fixed to 2 ms. The data was acquired with Compass otofControl (version 6.2, Bruker). The MS spectra and ion mobilograms were processed and integrated using DataAnalysis (version 4.0, Bruker). The MS spectra deconvolution was achieved with UniDec [^17^] (version 4.4.1), using a charge range of 1-20, molecular weight (MW) range of 1-20 kDa and a mass sampling every 10 kDa.

### Spin labeling and CW EPR spectroscopy

For spin labeling purified B-box protein (concentration 20 - 40 µM in 25 mM Tris-HCl pH 6.92, 150 mM NaCl containing buffer) was mixed with a stock solution of MTSL (TRC, Canada, cat. № TRC-O875000) spin label in acetonitrile (concentration: 20 mM) to obtain a molar MTSL to protein ratio of 1:0.9. The mixture was kept in the dark for 1 hour at room temperature for the spin labeling reaction to proceed.

The labeling efficiency and impact on the structure and conformation of the unfiltered protein solution was estimated by trapped ion mobility-mass spectrometry (TIM-MS) under native conditions (Figure S0). The deconvoluted MS spectra for the pristine and spin labeled protein shown in Figure S0 A and B, respectively, exhibit signals at 11460 Da (theoretical mass: 11453 Da) and 11640 Da which match the expected mass increase of 185 Da after spin labeling the single cysteine residue with MTSL. Based on the relative intensity in Figure S0 B a spin labeling efficiency close to 100% can be deduced. The IMS experiments add structural information on the labeled protein. This technique separates ions according to their size, shape, and charge. The collision cross-section (CCS) extracted from IMS corresponds to the rotationally-averaged surface of collision with the mobility gas. The IMS data of B-Box show that the principal contribution, at 1376 Å^2^ at charge state 6+ and between 1475 to 1517 Å^2^ at charge state 7+, is maintained after labeling. The data also show a distribution of higher CCS, around 1736 Å^2^ at charge state 7+ and 1477 Å^2^ at charge state 6+, which correspond to a more extended structure, i.e. unfolded protein. This partially unfolded contribution is present before labelling and not after labelling. Most of the pristine B-Box is thus found in a structured conformation, and a minor proportion is partially unfolded. The IMS mobilograms of the labelled B-Box display more homogeneous profiles, that are less altered by the contribution of the partially unfolded state. Overall, two protein variants are almost undistinguishable from one another.

Continuous wave (CW) EPR spectroscopy measurements were taken on a Bruker BR420 X-Band spectrometer upgraded with a Bruker ECS 041XG microwave bridge and a lock-in amplifier (Bruker ER023M) using a Bruker ER422 SHQ 8304 (SHQ) and 4119HS-W1/1136 (HQ) resonators at operating at frequencies around 9.862 GHz and 9.785 GHz, respectively; a modulation amplitude of 3 G and a modulation frequency of 100 kHz were used. The spectra were recorded with a receiver gain of 1e5, a time constant of 20.48 ms and a conversion time of 80 ms. The microwave attenuation was 20 dB. For measurements within the SHQ resonator the samples were placed in customized aqueous EPR flat cell optimized for the resonator (Hellma, Müllheim, Germany). 10 cm-long tubes with a 1 mm inner diameter (QSIL, Germany) were used for measurements in the HQ resonator. To minimize exposure to the electric field, the tube was filled up to approximately 1 cm of its length.

### AlphaFold modeling

ColabFold v1.5.5 [^18^] was used as an interface to access the AlphaFold-Multimer [^19^] plugin for AlphaFold2 [^20^] to predicted models for B-box multimers based on the sequence of the B-box domain. Default settings were used which results in five models for each monomer/oligomers studied. For each of these systems the structural model with the highest iDDT values was subsequently optimized to account for the ionic strength of the EPR buffer (150 mM NaCl) and a temperature of 298 K using the adaptive Poisson Boltzmann solver within CHARMM.

## Results and Discussion

### Formation of B-box associates during the expression procedure

The red trace in Figure 1 shows the dynamic light scattering (DLS) result of the as-prepared B-box sample recombinantly produced in E-coli. The experiment evidence the presence of larger agglomerates with a hydrodynamic radius around 32 nm. In addition, a broad peak around 2.3 nm is observed. This is still larger than the hydrodynamic radius expected for the monomeric form of the B-box. The observed tendency of the B-box to associate is expected based on the hydrophobicity plot [^21^] of HMGB1 revealing several hydrophobic patches (see SI, figure S2) which may cause the observed self-association behavior. To separate monomeric protein from the associated ones, the protein sample analyzed above was filtered through a membrane with a 30 kDa cutoff. This process resulted in a drastic change in the observed DLS result (see Figure 1, dark green line). After filtration a single narrow peak around 1 nm is observed which is well in-line with the expectation based on the size of the monomeric B-box showing that the procedure effectively removed all associated molecules from the solution [^22^]. The evaluation of UV-Vis intensity at 280 nm for the initial and filtered solutions (see SI, figure S4) revealed that approximately half of the protein molecules were present in the form of complexes. For the subsequent analysis of the structural and functional properties it is key to assess the stability of the monomers as it is well-known that some proteins can dynamically form functional multimers in solution [^23^]. Therefore, a series of DLS measurements were performed in 30-minutes intervals. No change in light scattering was observed over a 6-hour period, indicating that the monomeric protein is stable in solution over sufficient time that comparative characterization with other techniques is possible. It is worth mentioning that the A-box domain of the protein, which exhibits a reduced amount of hydrophobic patches, does not exhibit such association behavior (see SI Figure S3), which further suggests that hydrophobic interactions are involved in the formation of associates. Based on the observed stability of the monomeric form in solution, it is expected that the association may already occur in the molecularly crowded environment inside *E*.*Coli* cells during expression. As the latter condition might be closer to the physiologically relevant conditions than the much better defined monomeric form of the protein typically used for a detailed biophysical characterization it is of interest to elucidate the impact of the association not only on the structure of the protein but also on the functional properties such as the interaction with polyelectrolytes.

**Figure 1.**
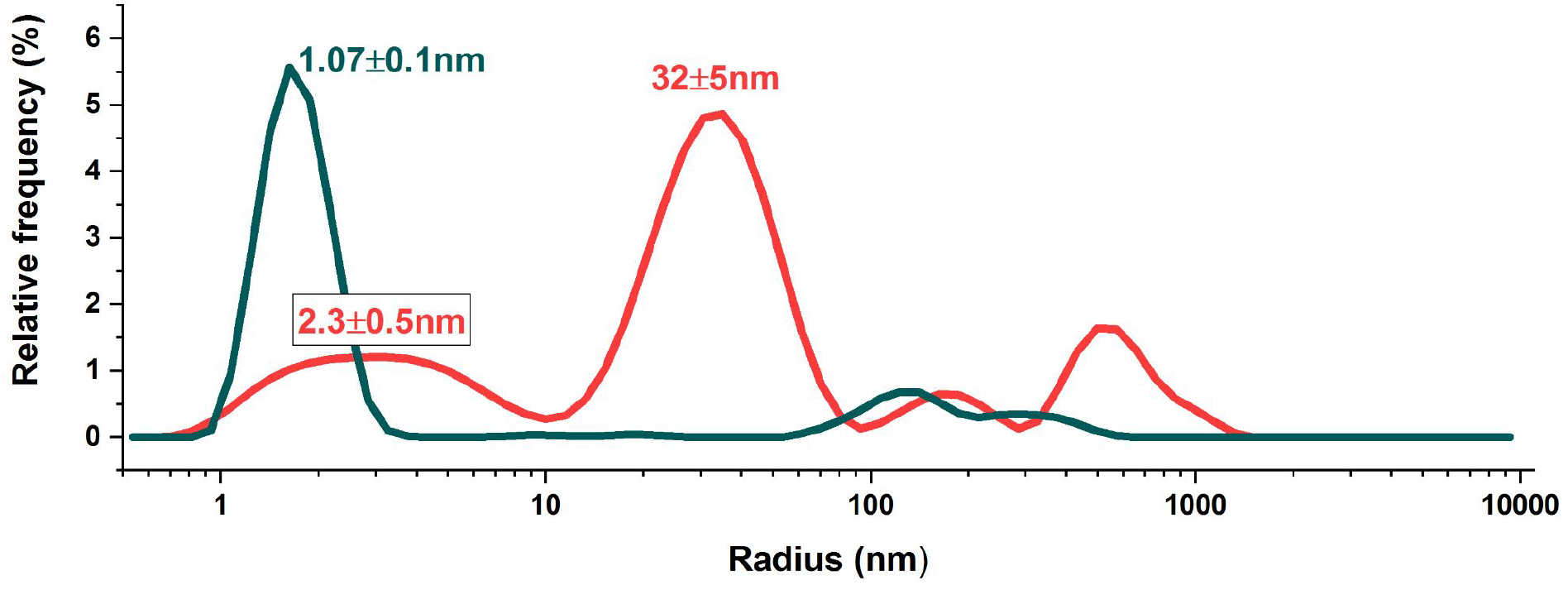
Hydrodynamic radius (R_h_) of B-box protein domain in buffer solution (25 mM Tris-HCl pH 6.9, 150 mM NaCl). Green line corresponds to B-box sample before filtration, where the presence of large multimeric complexes is evident, red line corresponds to a monomeric B-box sample filtered through cellulose membrane with 30 kDa cutoff filter.

### Impact of association on the structure of the B-box

To elucidate structural changes due to the association of protein, samples containing just monomeric species and those containing associated molecules were compared. Three different spectroscopic techniques which provide complementary insight were employed. Integral information on the secondary structure of the B-box was obtained from CD spectroscopy (see Figure 2A). The CD spectra of both species show very comparable spectra, with minima at 208 nm and 222 nm typical for α-helical proteins and consistent with the expectation based on the known crystal structure of the B-box [^24^]. These results clearly show that the secondary structure of the protein is very comparable for both samples. Complementary information can be obtained from fluorescence spectroscopy. The fluorescence of aromatic side chains were compared for both samples (Figure 2B) providing insight into the local environment of the corresponding side chains. Except for a reduced intensity observed for the associated state (red trace) both spectra are very similar, too. The fluorescence in this spectral range strongly depends on the properties of tryptophan residues. The B-box has only one tryptophan residue (Trp46) whose fluorescence is typically found in the range of 330 – 350 nm and four tyrosine residues whose fluorescence can contribute to the lower wavelength fluorescence (maximum around 305 nm). Due to the lower relative sensitivity of tyrosine compared to tryptophan (about 1:5), the strong dependence of the fluorescence maximum of tryptophan on polarity, and the possibility of resonance energy transfer (Förster radius of the Tyr-Trp pair: 4-16 Å) their relative contributions are rather difficult to disentangle. However, the lack of changes between the two samples excludes significant changes in the tertiary fold of the protein as the latter would cause a change in the local polarity of the tryptophan residue which is buried in the interior of the helical fold (Figure 2D) which is known to cause a shift in the fluorescence spectrum [^25^]. The difference in intensities can have different origins. One cause can be quenching - an effect well-known to occur at high local concentrations [^26^] but also other effects such as a redistribution of charges around the molecules in the different state can contribute to reduction of fluorescence intensity [^27^].

**Figure 2.**
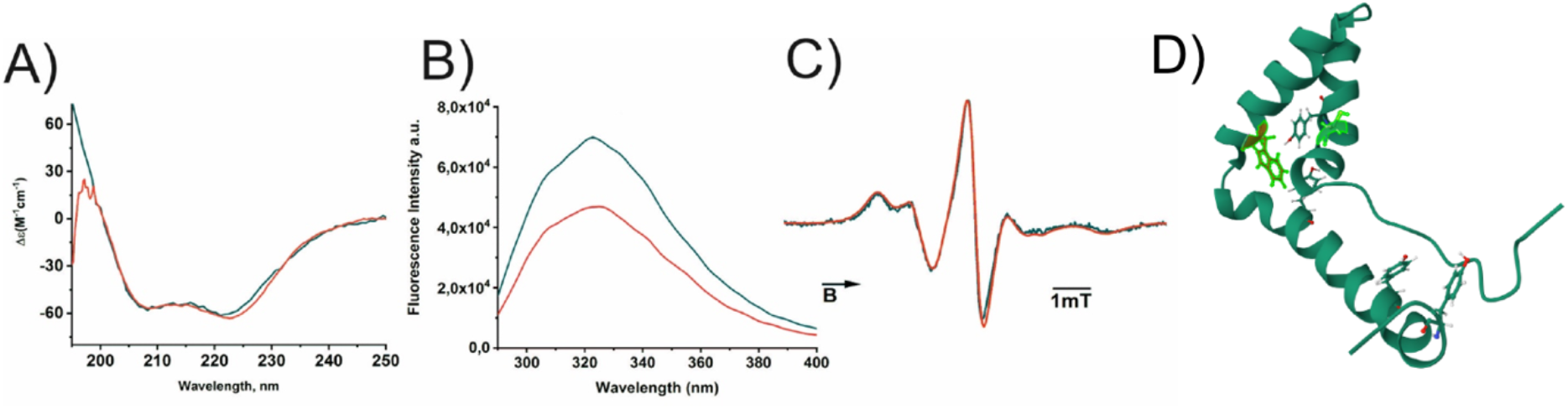
A) CD, B) fluorescence and C) EPR spectra of single (filtered through a membrane with 30 kDa cutoff) B-box protein (dark green) and unfiltered B-box protein (red) - where single protein molecules coexist with large self-complexes. D) structural model of the B-box with Trp and Cys residues highlighted (PDB: 1HMF).

As a third technique the EPR spectrum of an MTSL spin label attached to the only naturally occurring cysteine residue (Cys20; Cys106 in full length HMGB1) was analyzed [^28^]. According to the crystal structure Cys20 is a solvent exposed helical site. The EPR spectrum provides insight into the rotational dynamics of the side chain which can be altered by interaction in the vicinity of the spin label as well as a change of the global tumbling of the molecule [^29^]. The corresponding cw-EPR spectrum (Figure 2C (green line)) is perfectly in line with the expectation for a solvent exposed helical site. The spectrum for the sample containing associates (red spectrum Figure 2C) shows virtually the same spectrum, which further substantiate the conclusion drawn from the other methods. Moreover, the spectra provide no evidence for spin-spin interactions which excludes the presence of a significant portion of molecules with spin distances below about 1.5 nm [^30^].

Hence, all three techniques converge to the same conclusion i.e. the structure of the monomeric B-box is preserved even within the associated state. As the structure of the molecules are preserved these two systems provide the opportunity to study the differences in their functional properties e.g. their interaction with polyelectrolytes and allow to correlate possible changes in these properties with the local concentration of the protein as there is no interference with structural perturbation of the molecules due to the formation of the associates.

### B-box binds to heparin-sepharose

In 1989 Cardin *et al*. identified a consensus sequence of basic and non-basic amino acids which are found in several heparin binding proteins [^31^]. Two different motifs were determined as heparin-binding sequences, [-X-B-B-X-B-X-] and [-X-B-B-B-X-X-B-X-], in which B is a basic and X a hydropathic amino acid residue [^31^]. The motif (−P-K-K-P-R-G-K) present in the A-box of HMGB1 (residues 6-12) renders this domain the classical site for heparin binding. This is in line with surface plasmon resonance (SPR) studies which revealed low nanomolar affinity of HMGB1 to heparin and shows strong anti-inflammatory effects *in vitro* as well as *in vivo* by reducing the release of pro-inflammatory cytokines, like TNF-α, and blocking the interaction of HMGB1 to receptors on the cell surface [^32, 33^]. While the interaction of heparin with the consensus sequence within the A-box is expected, interaction of heparin with the B-box was not expected. However, the eluded fraction after the Ni-NTA affinity purification, which contains a protein with a molecular weight around 11 kDa-consistent with expectation for the B-box-already high purity, was found to bind to a heparin Sepharose column under low salt condition (50 mM NaCl) (Figure 4). This provides clear evidence for a binding of B-box motif of HMGB1 as expressed in E. Coli to heparin. As the B-box can be eluded from the heparin column with a high ionic strength buffer (2 M NaCl, trace 11, Figure 4) it can be concluded that the interaction has large electrostatic contribution. This conclusion is substantiated by a Western blot with a B-box specific α-HMGB1 antibody (detecting aa. 150-177 of the full-length protein) proving the identity of the purified B-box protein (Figure 4D). It is worth mentioning that SDS-PAGE provides no evidence for the existence of multimers observed in solution. This indicates that the denaturing conditions of the SDS-PAGE disrupt the associates, which is in line with expectations based on the weak perturbation of the protein structure within the associates which renders a weak interaction between the monomers likely.

**Figure 4:**
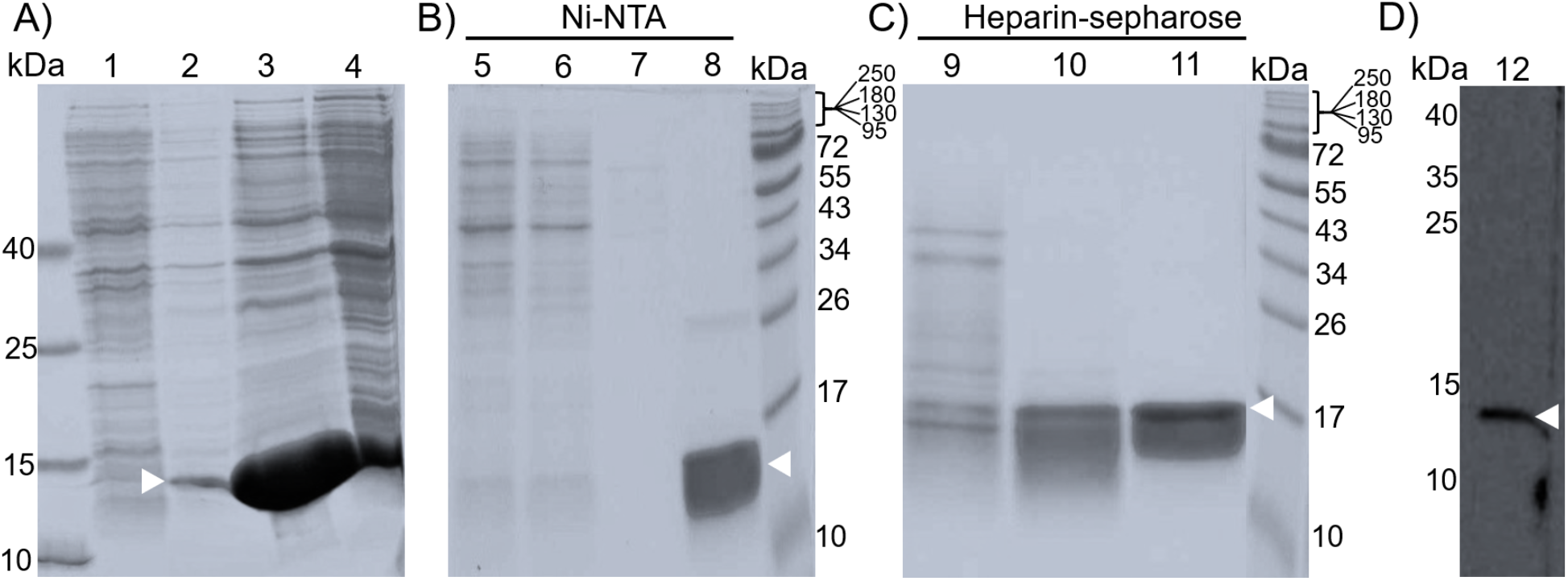
Analysis of expression and purification of the HMGB1 B-box. Samples of the expression and purification steps were analyzed on a 15%-SDS PAGE A) proof of soluble expression, B) purification via Ni-NTA agarose, C) purification via heparin sepharose and D) by Western blotting using 50 ng purified B-box and α-HMGB1 antibody detecting an epitope between amino acid 150-215. The protein migrates around 11.4 kDa, which is in line with the theoretical molecular mass. The white arrow indicates the corresponding band for the B-box. A) 1: fraction from non-induced culture, 2: fraction from induced culture, 3: soluble extract, 4: insoluble extract. B) 5: flow through Ni-NTA, 6: wash (50 mM Tris-HCl, pH 8.0, 1.5 M NaCl, 20 mM imidazol), 7: wash (50 mM Tris-HCl, pH 8.0, 50 mM NaCl, 20 mM imidazol), 8: elution (50 mM Tris-HCl, pH 8.0, 50 mM NaCl, 500 mM imidazole. C) 9: flow through heparin column, 10: wash (50 mM Tris-HCl, pH 8.0, 100 mM NaCl), 11: elution (50 mM Tris-HCl, pH 8.0, 2 M NaCl). D) 12: Blot with B-box specific α-HMGB1 antibody.

### Heparin binds B-box in solution

Next, we tested if the B-box complexes also bind heparin in solution. Therefore, we incubated FITC labeled heparin with the protein in equal amounts (10 µg each) in a buffer containing 50 mM Tris-HCl pH 8.0, 150 mM NaCl and analyzed the samples on a native PAGE. As could be seen in Figure 5 (lane 1), the heparin-FITC sample without the B-box runs as a heterogenous band with a strong signal in the low molecular range. This outcome can be expected from heparin, as polymer samples generally exhibit a significant molecular mass distribution even for a fractionated heparin sample with a molecular weight of 15 kDa, used here. While the B-box itself gives no fluorescent signal (Figure 5, lane 2), an incubation with the heparin-FITC results in band corresponding to a slower migrating species (Figure 5, lane 3) which shows the formation of species with higher molecular weight expected for the complex between heparin and the B-box. The incomplete binding of heparin-FITC to the B-box (broad band at low molecular weight, Figure 5, lane 3) could be due to a lower binding affinity or an overload of heparin. The Coomassie stained gel also showed a complex of heparin-FITC and B-box at the same height as could be seen in the FITC visualized gel (Figure 5, lane 3). Nevertheless, this experiment further shows that the B-box in the complex form is capable of binding heparin in solution and builds a stable complex that can be separated by a native PAGE.

**Figure 5:**
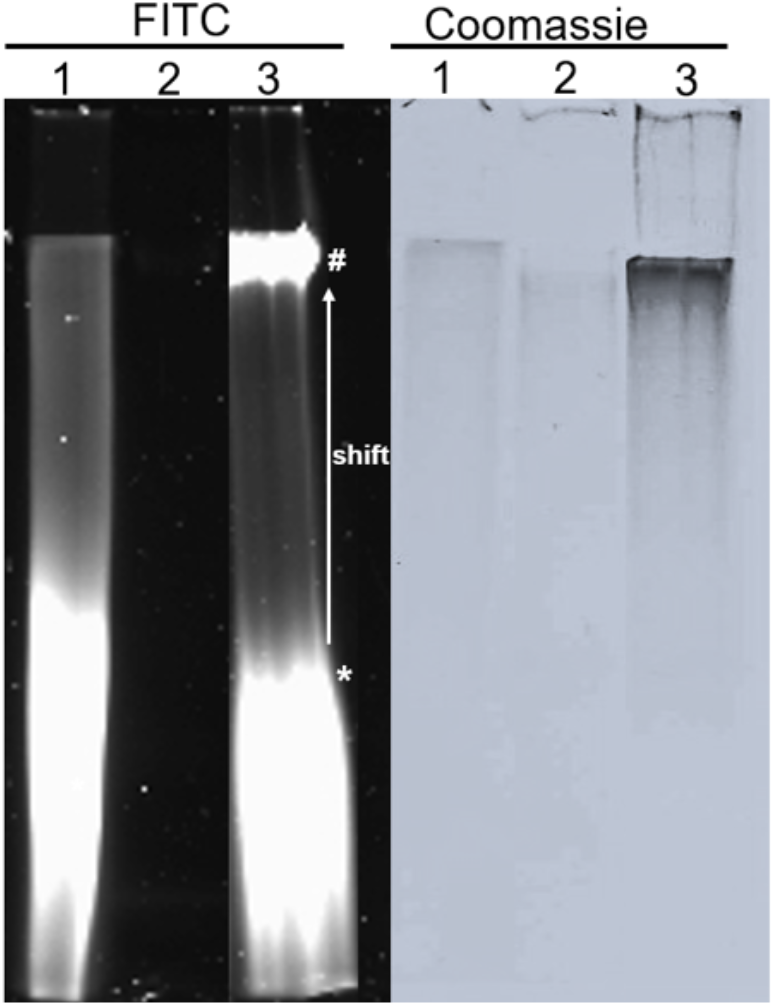
B-box binds Heparin. FITC labeled heparin was incubated with the B-box domain in equal amounts of 10 µg. Left: complex formation was confirmed by 12% native PAGE and imaging in the FITC channel (absorption 501 nm, emission 523 nm). Right: Coomassie blue staining. 1, 10 µg heparin-FITC, 2, 10 µg B-box, 3, heparin-FITC + B-box. * heparin-FITC. # complex heparin-FITC and B-box.

### B-box binding patterns to heparin in single and self-associated states

The results presented above revealed that the B-box in self-associated state can bind to heparin. EPR spectra of the spin labeled B-box (MTSL at Cys20) can provide access to the impact of heparin binding on the structure of the B-box. Figure 6 shows the EPR spectra of the B-box in the monomeric (a) and the associated (b) state prior (black traces) and after adding different amounts (red trace 15 mol%; blue traces 100 mol%) of heparin.

**Figure 6:**
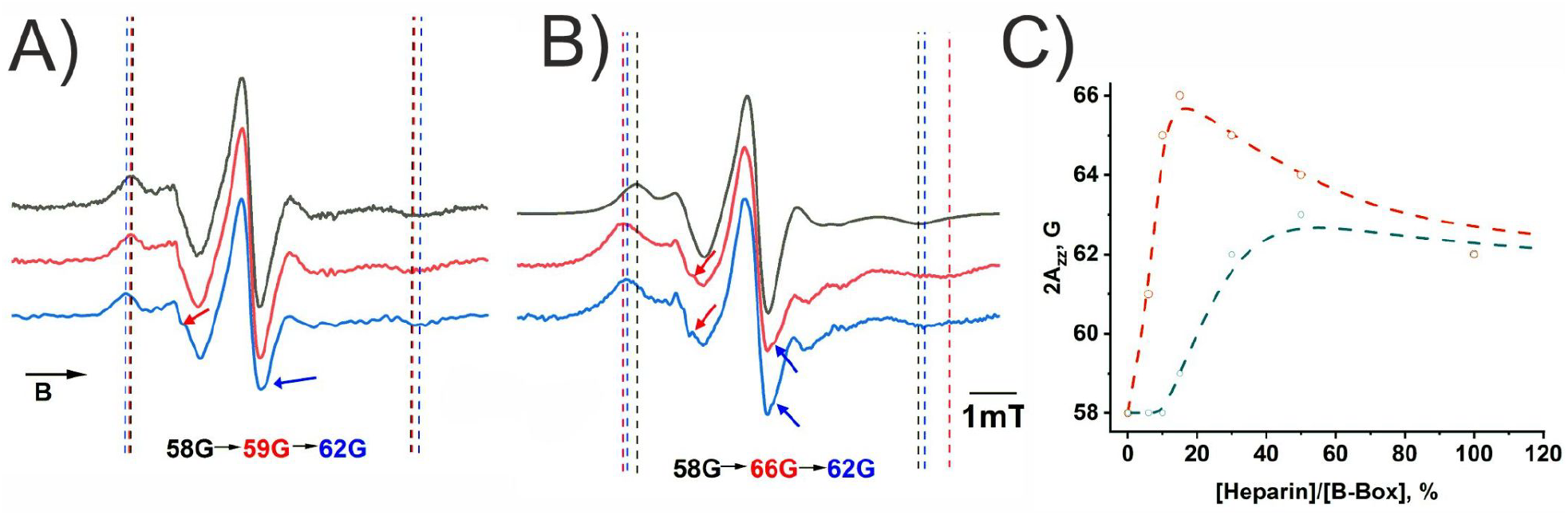
EPR spectra of samples containing a) monomeric and b) associated B-box proteins prior to (black lines) and after the addition of 15 mol% (red lines) and 100 mol% (blue lines) of heparin. c) splitting between the outer exprema (2A_zz_) as a function of added heparin (mol%). For the corresponding set of spectra, see SI Figure S7.

As seen from Figure 6A the addition of 15 mol% heparin to the B-box has little influence on the splitting between the outer extrema of the spectra corresponding to 2A_zz_. The latter value starts to change only after addition of approximately 20 mol% of heparin (Figure 6C). The increase of A_zz_ indicates changes in the rotational dynamics of the spin label which is associated to the interaction of the B-box with heparin. This result is consistent with a low binding affinity of the monomeric B-box to heparin as expected due to the lack of a consensus binding sequence for heparin. In contrast, for the B-box sample containing associated protein molecules shows a notable increase in the maximum splitting value already after adding 6 mol% of heparin (Figure 6C). This suggests that the self-association of the B-box protein increases the interaction strength to the negatively charged polyelectrolyte heparin considerably. An important question concerns the origin of the experimentally observed variation in the binding affinity of the B-box to the polyelectrolyte. A likely origin of the interaction is electrostatics which poses the question as to how this interaction is altered in associated state as compared to the monomeric protein. To this end, we employed Alphafold2 to predict structures for a series of oligomers of the B-box. Figure 7 shows the structure for the most likely structure for oligomers containing between two and eight monomeric units together with the corresponding charge distribution calculated for the respective structures.

**Figure 7:**
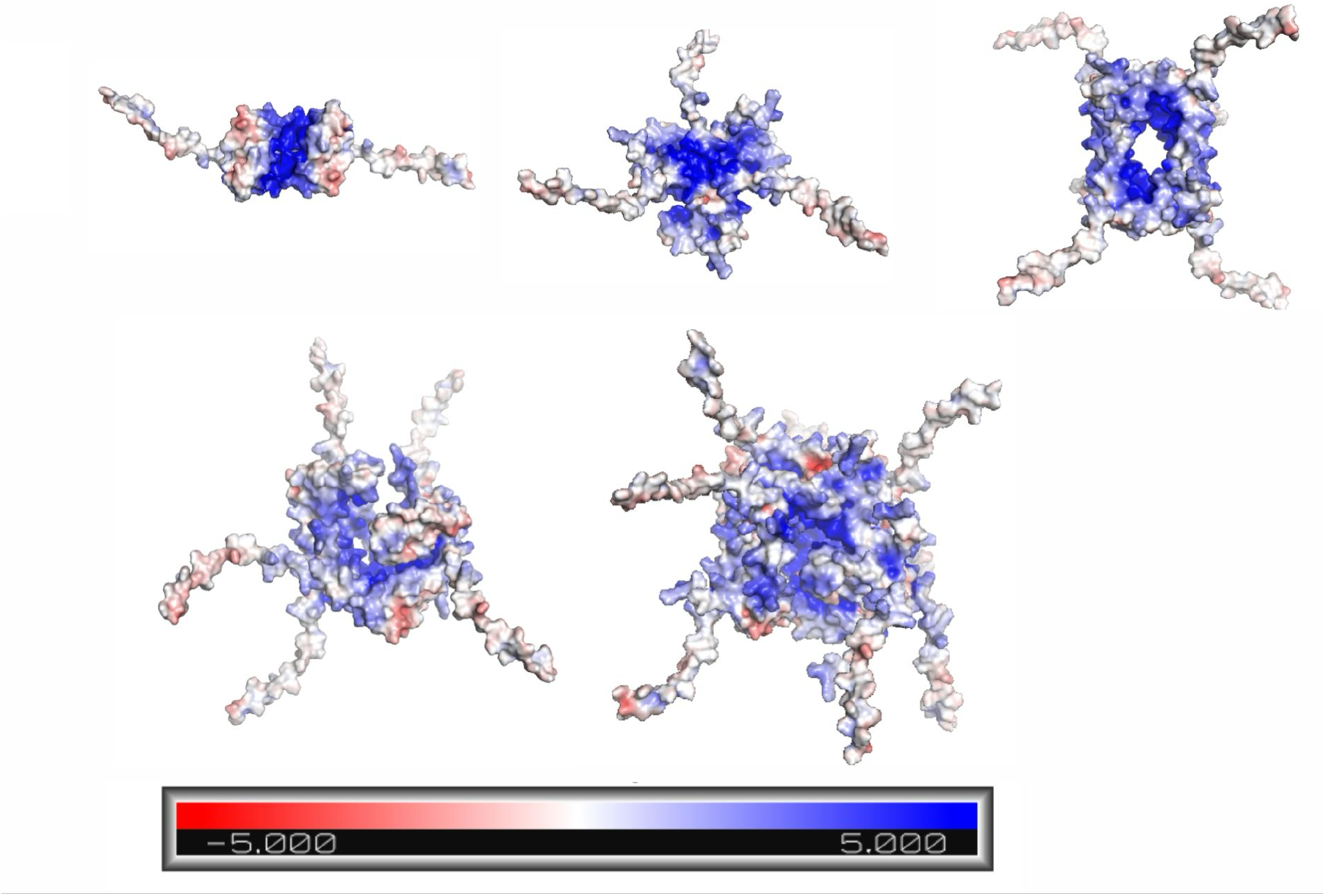
Charge distribution models of AlphaFold2 predictions for B-box monomer, dimer, trimer, tetramer, pentamer and octamer, with each panel representing an increase in multimer complexity respectively.

As the number of B-box units increases, the simulations predict patches of positive charge (blue color) in the central region of the structural models. Please note that the oligomers modeled here are not meant to serve as structural models for the associates present in the B-box sample investigated by EPR spectroscopy as the observed size of the associates (32 nm) requires much larger number of protein molecules. However, as the oligomers modelled here consistently show patches of positive charges independent of their size this property can be considered a common feature of the associates also found for larger sizes. In turn, this agglomeration of positive charge whose surface area increases with size is likely to increase the strength of electrostatic interaction to negatively charged species such as heparin and can hence provide a rationale for the observed increased interaction strength of the B-box with heparin.

Interesting additional structural information on the impact of heparin binding to the B-box can be obtained from an analysis of the EPR line shape. Comparing the values of 2A_zz_ for the two samples as a function of heparin concentration considerable differences can be observed beyond the different onset discussed above. The associated form of the B-box exhibits an increase of the maximal splitting by about 8 G after adding 15 mol% heparin which decreases upon further addition of heparin to about of that value (4 G, 2A_zz_ = 62 G) after a large excess of heparin (400 mol% heparin; s. Figure S7). Importantly, an identical splitting is observed for the monomeric solution with the same molar ratio of heparin (Figure S7) which indicates that the final state after adding excess of heparin is comparable for both systems. In this respect it is also important to mention that in both cases the final spectra as well as those with equal molar amount of heparin exhibit additional spectral changes as indicated by the arrows in Figure 6A and B. The broadening of the high-field minimum of the central line (blue arrows) is consistent with a reduced averaging of the z-component of the g-tensor. The occurrence of resolved z-components is typically associated with a reduced amplitude of the anisotropic motion of the spin label i.e. a higher order parameter of the z-axis. Such changes are typically observed for buried sites or spin labels in which the amplitude is structurally restricted such as a bidentate label [^34, 35, 36^]. In the present case the occurrence of this reduction in amplitude provides direct experimental evidence that the interaction site of the heparin is close to the Cys 20 residue of the B-box. It is interesting to note that this change is observed for both samples at high heparin concentration indicating that the interaction site of heparin is affecting the spin label dynamics similarly in both samples consistent with comparable interaction sites. For the sample containing associated molecules the corresponding changes are observed already for small heparin concentration (s. red spectrum in Figure 6B). In this respect it is important that the restriction of the local motion of the spin label remains even though the overall width of the spectrum decreases (compare red and blue trace in Figure 6B and red symbols in Figure 6C). The latter is interpreted as being due to an increase of global rotational correlation time of the B-box/heparin complexes due to a reduction in their size with increasing heparin concentration. Based on the similarity of the spectral width and line shape observed for both samples for an excess of heparin both samples are expected to exhibit a comparable size of the heparin/B-box complexes in the limit of large heparin excess.

## Conclusions

The domains of the HMGB1 protein are known to mediate interactions with distinct molecules, including various polyelectrolytes, as well as the process of liquid-liquid phase separation in crowded cellular environments. Traditionally, it has been believed that only the N-terminus of the A-box provides the binding affinity to the highly negatively charged polyelectrolyte, heparin. However, our investigation demonstrates that the B-box domain also binds to heparin, and its binding affinity is regulated by the formation of dynamic self-complexes between B-box proteins. We found that the self-association of the B-box despite the structural similarity of the protein in monomeric and associated state increases the binding affinity to polyelectrolytes considerably. It was shown that oligomerization can lead to an accumulation of positive charge in the core of the oligomers which provides a rationale for the increased interaction strength to negatively charged heparin. EPR spectroscopy provide evidence for an interaction site between the B-box and heparin close to the Cys20 residue and also suggests that the interaction with heparin leads to changes of the association of the B-box. Above equal molarity of B-box and heparin both samples exhibit similar EPR spectra which not only suggest comparable interaction sites but also similar overall molecular weight of the heparin/B-box complexes in the limit of high heparin excess.

## Acknowledgments

This work was funded by the Deutsche Forschungsgemeinschaft (DFG, German Research Foundation) – 434130070 within the GRK 2662.

