## Supplementary Material for "HMGB1 B-box domain self-complexes promote protein-polyelectrolyte interactions"

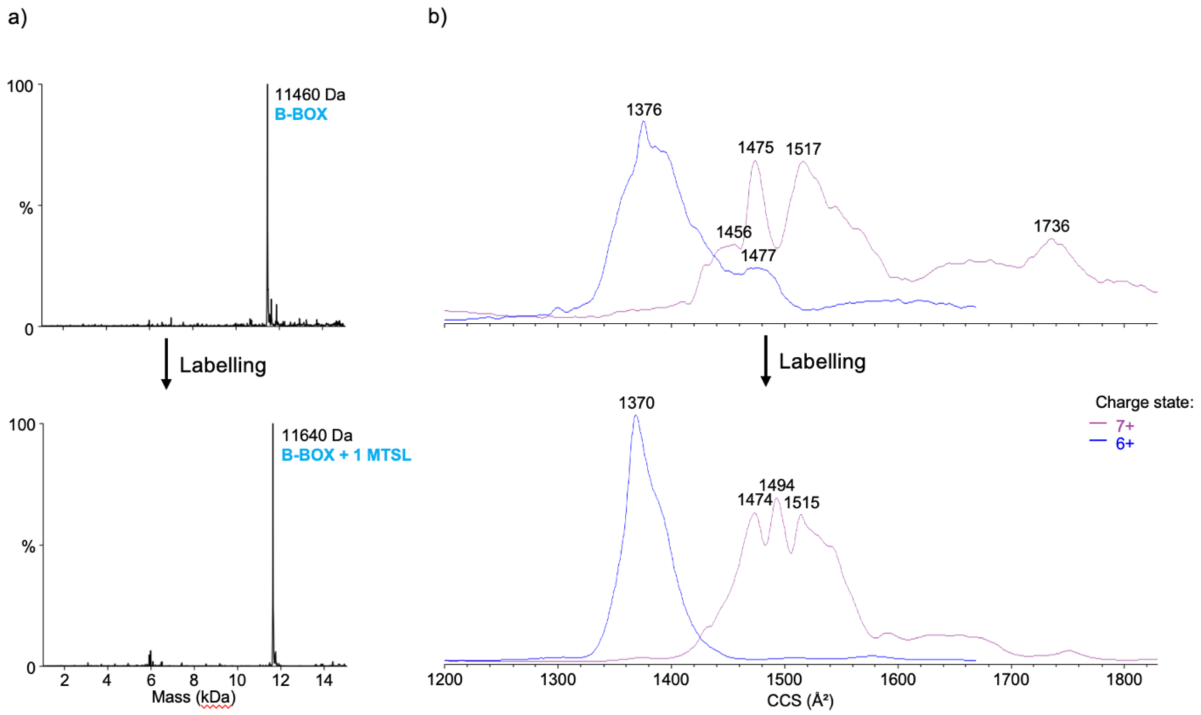


**Figure S0.** a) Native ion mobility-mass spectrometry (IM-MS) experiments on the unfiltered B-box domain before and after MTSL labeling. a) Deconvoluted mass spectra and b) charge distribution plots.

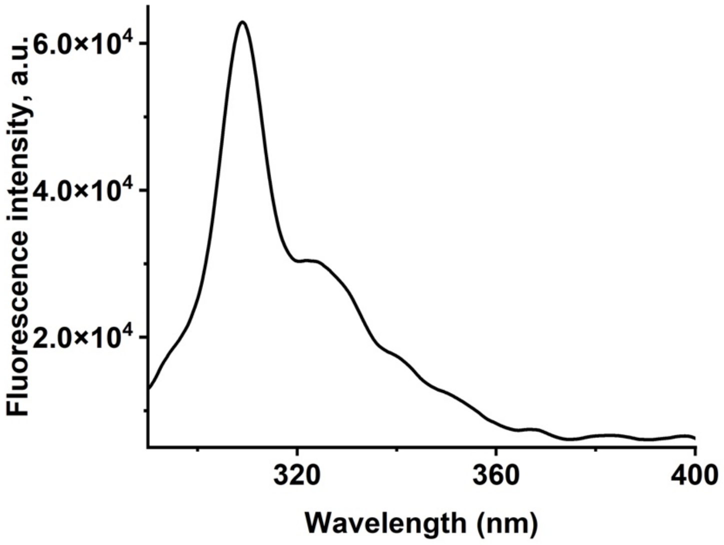


**Figure S1.** Baseline (buffer solution, 25 mM Tris, 150 mM NaCl, pH 6.9) for the fluorescence experiments. The Raman scattering peak is visible at around 310 nm.

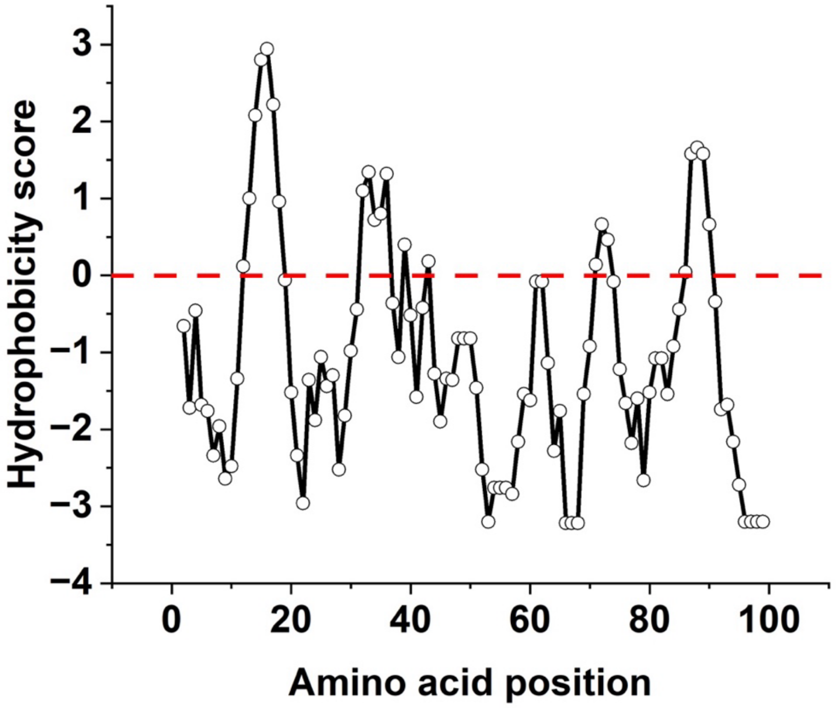


**Figure S2.** Hydrophobicity of the B-Box domain along the chain.

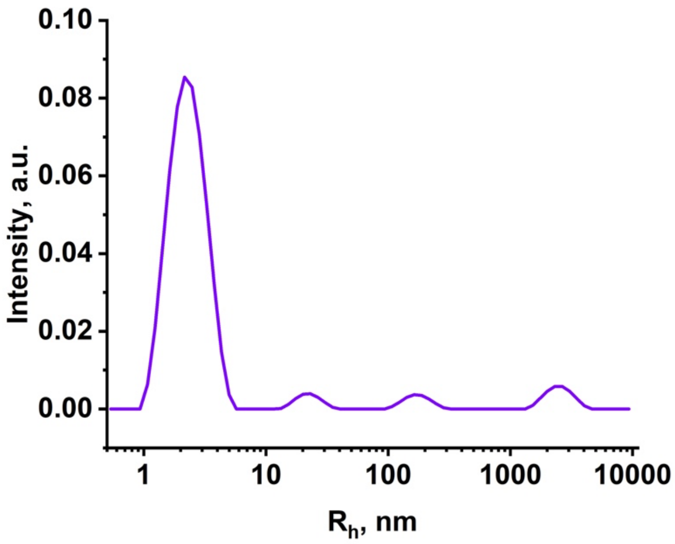


**Figure S3.** Hydrodynamic radius ($R_h$) of A-box HMGB1 protein domain in buffer solution (25 mM Tris-HCl pH 6.9, 150 mM NaCl).

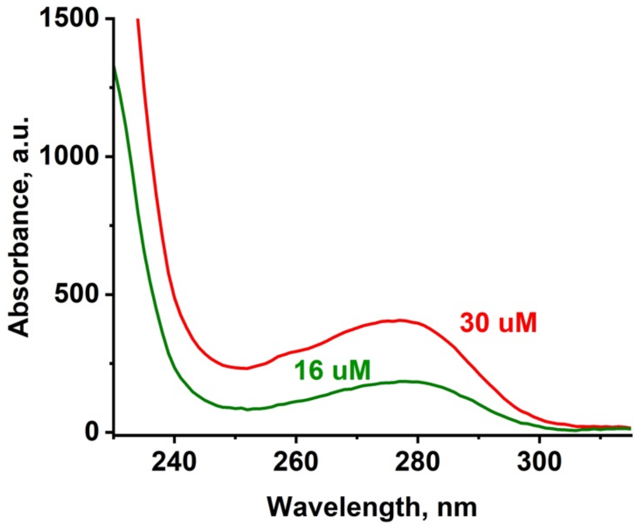


**Figure S4.** UV-vis spectra of the B-Box solution before (red line) and after (dark green line) the filtration through 30 kDa membrane filter and corresponding concentration measured according to the intensity at 280 nm.

To perform binding studies, we cloned the B-box into a modified variant of the pet22b(+) vector that includes a C-terminal linker region (S-G-S) and two additional histidine residues (8 histidines in total) for better purification. To obtain a stable expression we also added the amino acid triplet A-R-I to the N-terminus [14]. The results of the cloning procedure can be found in Supplementary, Figure S5, S6. The expression of recombinant human B-box protein was done with BL21(DE3) *E. coli* cells and the protein was highly expressed in a soluble form, as could be seen in Figure 4.

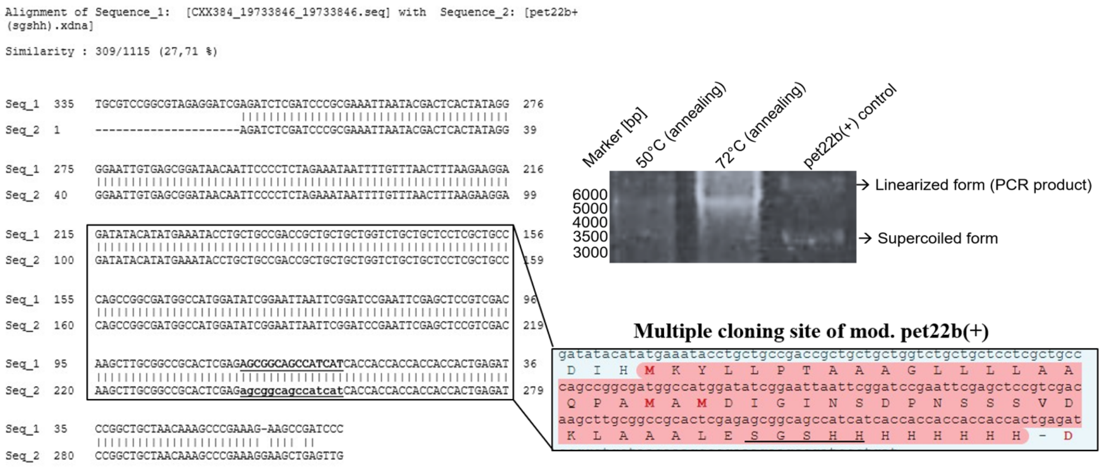


**Figure S5.** Cloning results of the C-terminal modification of the pet22 b(+) vector. The C-terminal insertion of a Serine-Glycine-Serine (SGS) and two histidine´s was achieved by PCR and blunt end cloning. Successful cloning was verified by sequencing and the inserted amino acids are underlined.

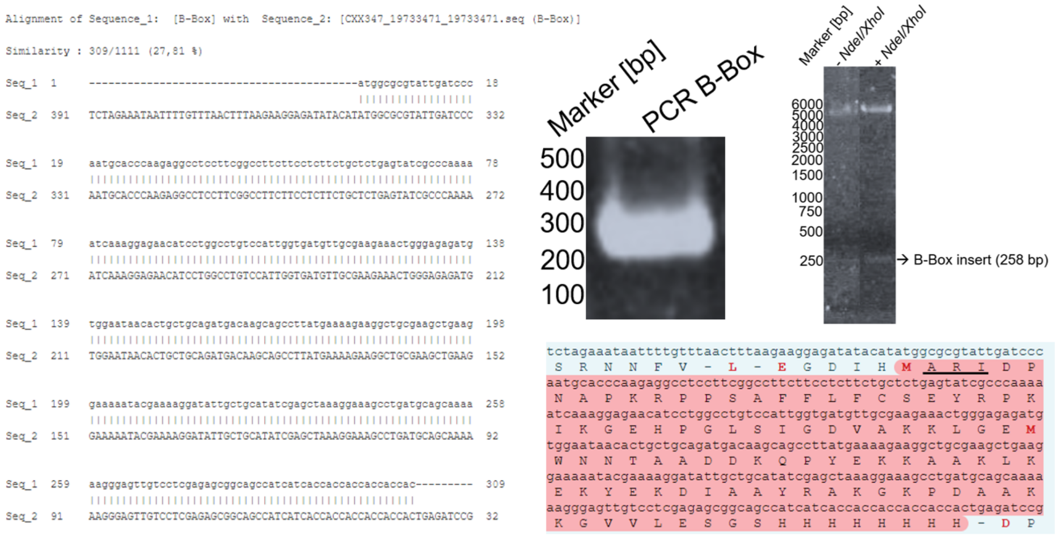


**Figure S6.** Cloning of human B-box into mod. pet22 b(+) vector. The sequence of the human B-box protein from HMGB1 was cloned into the mod. pet22 b(+) vector by ELIC. Successful cloning was verified by sequencing and the corresponding amino acid sequence was highlighted in pink. The three artifical amino acids Alanin-Arginin-Isoleucin are underlined.

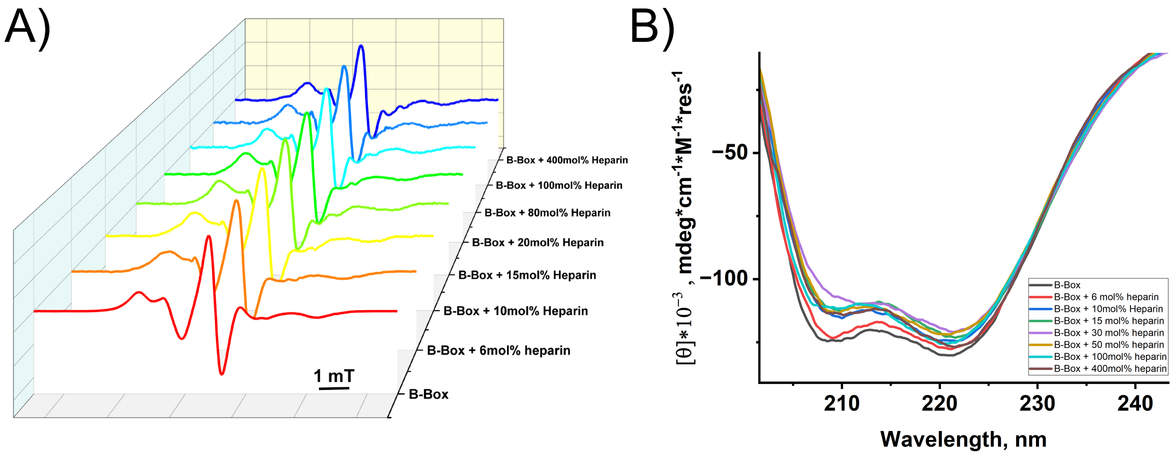


**Figure S7.** A) EPR spectra of HMGB1 B-box domain protein after gradual addition of heparin. B) CD spectra showing the influence of different concentrations of heparin (mol%) on the HMGB1 B-Box secondary structure.
